# AFNI and Clustering: False Positive Rates Redux

**DOI:** 10.1101/065862

**Authors:** Robert W Cox, Richard C Reynolds, Paul A Taylor

## Abstract

In response to reports of inflated false positive rate (FPR) in FMRI group analysis tools, a series of replications, investigations, and software modifications were made to address this issue. While these investigations continue, significant progress has been made to adapt AFNI to fix such problems. Two separate lines of changes have been made. First, a long-tailed model for the spatial correlation of the FMRI noise characterized by autocorrelation function (ACF) was developed and implemented into the 3dClustSim tool for determining the cluster-size threshold to use for a given voxel-wise threshold. Second, the 3dttest++ program was modified to do randomization of the voxel-wise t-tests and then to feed those randomized *t*-statistic maps into 3dClustSim directly for cluster-size threshold determination-without any spatial model for the ACF. These approaches were tested with the Beijing subset of the FCON-1000 data collection. The first approach shows markedly improved (reduced) FPR, but in many cases is still above the nominal 5%. The second approach shows FPRs clustered tightly about 5% across all per-voxel *p*-value thresholds ≤ 0.01. If *t*-tests from a univariate GLM are adequate for the group analysis in question, the second approach is what the AFNI group currently recommends for thresholding. If more complex per-voxel statistical analyses are required (where permutation/randomization is impracticable), then our current recommendation is to use the new ACF modeling approach coupled with a per-voxel *p*-threshold of 0.001 or below. Simulations were also repeated with the now infamously “buggy” version of 3dClustSim: the effect of the bug on FPRs was minimal (of order a few percent).

## Introduction

The reports [1,2] of inflated false positive rates (FPRs) for commonly used cluster-threshold based FMRI statistics packages (SPM, FSL, AFNI) caused a stir in technical and semi-popular publications. The responses to the dramatic statement ([2]; since mollified [3]) that, “These results question the validity of some 40,000 FMRI studies,” have ranged from saying that there is nothing new here (“tests based upon spatial extent become inexact at low thresholds” [4]), to cautious-but-concerned commentary [5], to hyperbole about invalidating 15 years of research due to a software bug [6,7].

The AFNI team takes the question of the inflated FPRs seriously, but does not consider that the FMRI apocalypse has arrived. In particular, since the specific “bug” referred to in [2,6,7] was in the AFNI program 3dClustSim, we feel it necessary to report specifically on the changes made to the AFNI package to adapt to these problems.

This report is divided into two themes. First, looking to the past, we show the effect of the erstwhile bug within 3dClustSim (non-negligible, but non-monstrous). Secondly, looking forward, we present two separate approaches that have been taken to rein in the FPRs, both of whose development is ongoing and which were first presented at the 2016 OHBM meeting [8]. The first is to refine the statistical model underlying the parametric model for calculating cluster-size thresholds for a given per-voxel *p*-value threshold. The second approach is to eschew a direct model and instead use a randomization approach to generate cluster-size thresholds from the residuals of the inter-subject (group) *t*-test. Sets of simulations following those in [2] are presented.

## Methods: Simulations

We repeated the simulation steps carried out in [1,2] using the 198 Beijing datasets from the FCON-1000 collection [9]. The detailed AFNI processing for each subject was somewhat different than used in [2], since we ran with our most up-to-date recommendations for time series analyses (eg, using AnatICOR and nonlinear registration; see afni_proc.py’s helpfile Example 11), but the results are quite comparable. The group analyses, using 1000 random sub-collections of the Beijing datasets, were carried out using 3dttest++ in the same way as described in [2]; in particular, using exactly the same sets of sub-collections (made possible by the authors of [2] putting their processing scripts on GitHub).

In the initial work [2], various combinations of simulation parameters produced widely varying levels of agreement or disagreement with the nominal 5% setting for FPRs for all software tools tested. In order to present a broad description, the comparisons presented here are for the full set of basic simulations put forth in [2]. This includes investigating the four values of Gaussian smoothing applied (Full-Width at Half-Maximum (FWHM) of 4, 6, 8 and 10 mm) and two different per-voxel *p*-value thresholds (0.01 and 0.001). Additionally, four separate stimulus timings were used: blocks of 10 s ON/OFF (“B1”) and 30 s ON/OFF (“B2”), and event-related paradigms of (regular) 2 s task with 6 s rest (“E1”) and (random) 1-4 s task with 6 s rest (“E2”).

## Results

We present results of the simulations re-run with the various changes to AFNI. We then investigate various problems that exist in cluster detection tools, their impacts on results (i.e., FPRs), and preliminary means for correcting/addressing them.

### The Past-3dClustSim and “The Bug.”

The first problem was particular to AFNI: there was a bug in 3dClustSim. This program works by generating a 3D grid of *N*(0,1) iid random deviates, then smoothing them to the level estimated from the residuals of the FMRI data model, then carrying out voxel-wise thresholding followed by clustering to determine the rate at which clusters of different sizes occur at the various voxel-wise thresholds. The bug, pointed out by the authors of [2] in an email, was a flaw in how 3dClustSim rescaled the 3D noise grid after smoothing in order to bring the variance of the values back to 1.0 (for ease of later *p*-value thresholding). This rescaling was off due to improper allowance for edge effects, with the result being that the cluster-size thresholds computed were slightly too small, so that the FPR would end up somewhat inflated. During part of the work leading to [1,2], this bug was fixed in May 2015, and noted in the log of AFNI software changes (https://afni.nimh.nih.gov/pub/dist/doc/program_help/history_all.html):

> 12 May 2015, RW Cox, 3dClustSim, level 2 (MINOR), type 5 (MODIFY) Eliminate edge effects of smoothing by padding and unpadding
>
> Simulate extra-size volumes then smooth, then cut back to the desired volume size. Can use new ‘-nopad’ option to try the old-fashioned method. (H/T to Anders Eklund and Tom Nichols.)

Results comparing the pre- and post-fix versions of the standard 3dClustSim (“buggy” and “fixed”, respectively) are shown in Fig. 1, which presents FPRs from re-running the two-sample *t*-tests (40 subjects total per each of 1000 3D tests) of [2]. The effects of the bug were modest, particularly for p=0.001 where the FPR increases were < 1-2%. At the less stringent per-voxel p=0.01, where FPRs had been noticeably more inflated for most software packages, the difference was greatest for the largest smoothing (understandably, given the problem within the program), approximately < 3-5%. In each case, the difference between the “buggy” and “fixed” values was very small compared to the estimated FPR, meaning the bug had only a minor impact. At *p*=0.001, the “fixed” results for the event-related stimulus timings are not far from the nominal 5% FPR (Fig. 1, lower panel); however, the corresponding “fixed” results for the block-design stimulus timings are still somewhat high. The p=0.01 results are all still far too high in the “fixed” column (Fig. 1, upper panel).

**Figure 1.**
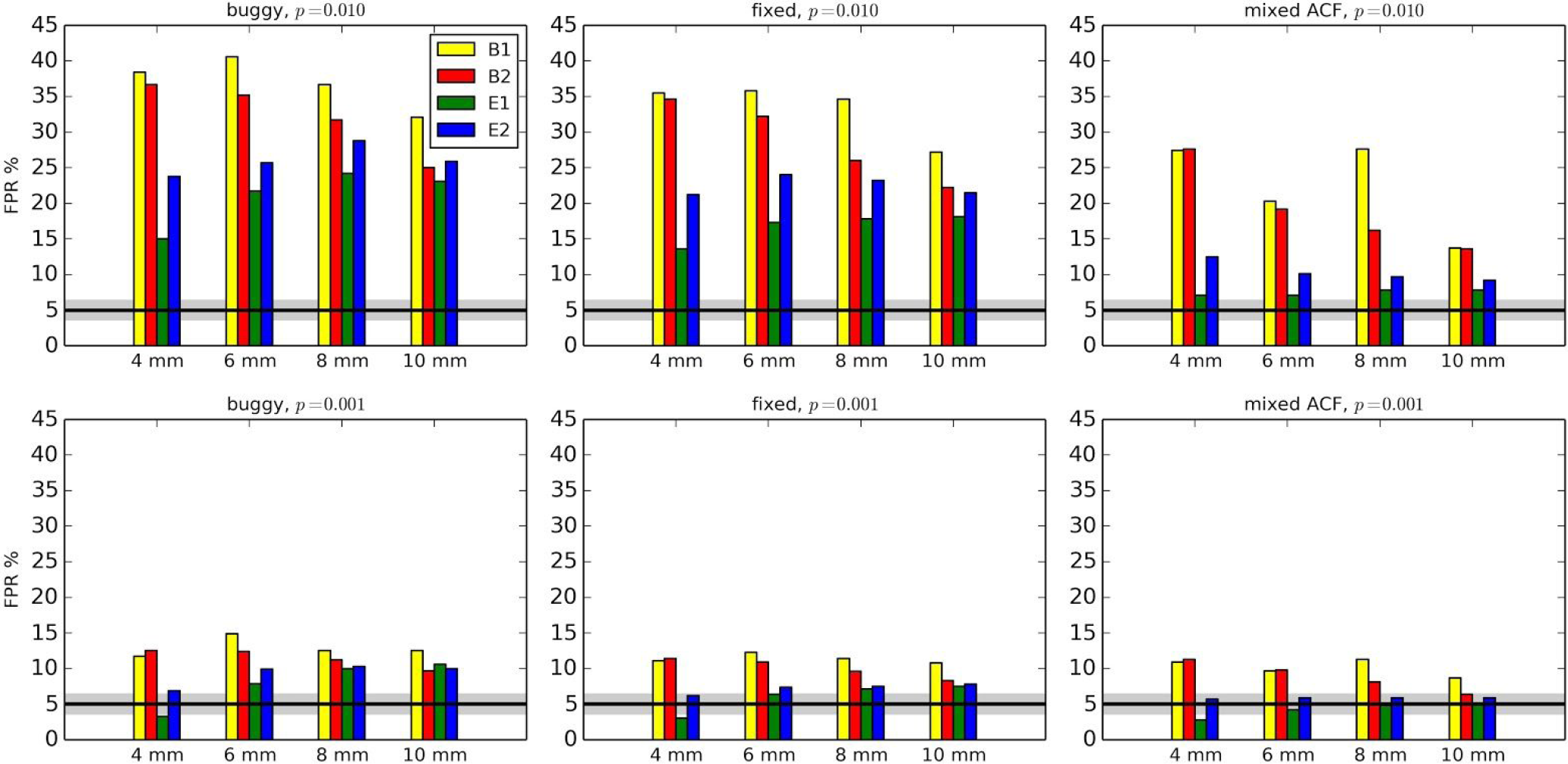
False Positive Rates (FPRs) for various software scenarios, with 1000 2-sample *t*-tests (as in [1,2]) using 20 subjects’ data in each sample. “buggy” and “fixed” means the cluster-size thresholds were selected using the Gaussian shape model with the FWHM being the median of the 40 individual subject’s values: “buggy” via 3dClustSim before the bug fix, “fixed” via 3dClustSim after the bug fix. “mixed ACF” means the cluster-size threshold was selected using the (Eq 1) model for spatial correlation of the noise, with the *a,b,c* parameters being the median of the 40 individual subject’s values (estimated via program 3dFWHMx). Two different per-voxel *p*-value thresholds (1-sided tests, as used in [2]) are shown. The black line shows the nominal 5% false positive rate (out of 1000 trials), and the gray band shows its theoretical 95% confidence interval, 3.65-6.35%. As in [2], different smoothing values were tested (4-10 mm). B1 = 10 s block; B2 = 30 s block; E1 = regular event related; E2 = randomized event related.

### The Present-Updating “The Flaw”-Assumptions about Spatial Smoothness

The second problem in determining cluster-size is much more widespread (to date) across the tools most used in the FMRI community: it is the flawed assumption that the shape of the spatial autocorrelation function (ACF) in the FMRI noise is Gaussian in form. That is, it is generally assumed that, for voxels separated by Euclidean distance *r*, the spatial correlation between noise values has the form exp[*-r^2^/(2b^2^)*], where it is traditional to specify the parameter *b* by the equivalent FWHM of [8 ln(2)]^1/2^×*b* = 2.35482×*b*. In fact (as pointed out in [2]), the empirical ACF shape, computed from the FMRI residuals and averaged across the whole brain, has much longer tails than the Gaussian shape. The heavy-tailed nature of spatial smoothness within the brain, which had been largely ignored previously, has significant consequences for thresholding clusters in FMRI analyses.

Fig. 2 illustrates the problem, along with the current solution adopted in AFNI. The empirical correlation falls off rapidly with *r* at first, but then tails off much slower than the Gaussian function. We found that the empirical ACF estimates are typically well fit by a function of the mixed Gaussian plus mono-exponential form

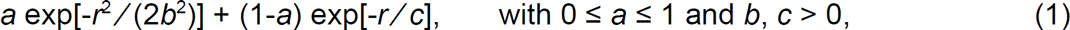

**Figure 2.**
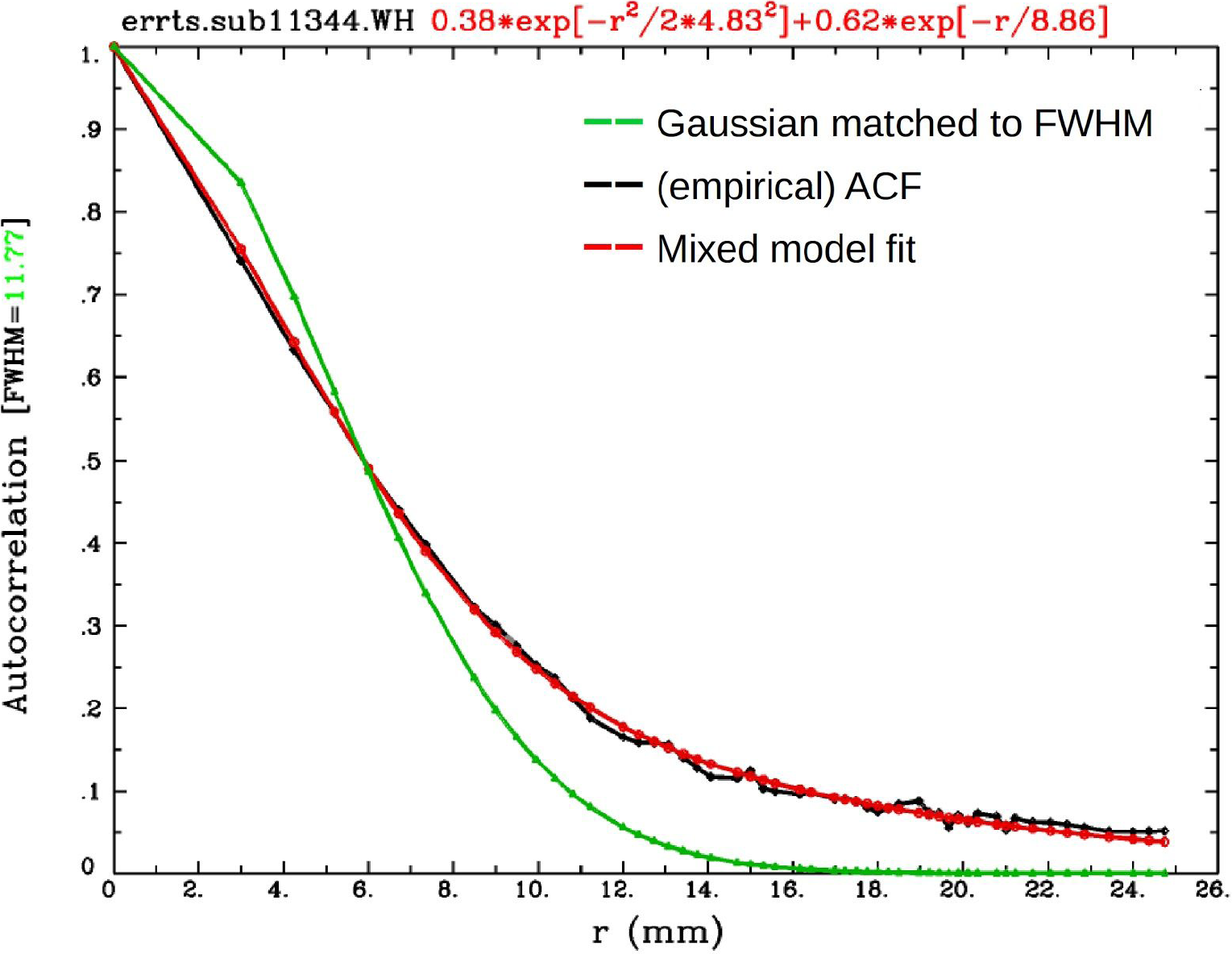
An example comparison of the empirical ACF (black) and the estimated Gaussian fit (green), which have large differences (importantly, in the tail drop-off above *r* ~ 8 mm). The proposed mixed model after fitting parameters as described in (Eq 1) provides a much better fit of the data.

which is illustrated in Fig. 2. Once the inadequacy of the pure Gaussian model was realized, 3dClustSim was modified to allow the generation of random 3D fields with autocorrelation given by (Eq 1), using a FFT approach (which does not lead to prohibitive runtimes). FPRs from re-running the 2-sample *t*-tests of [2], using the (hopefully bug-free) ACF model option in 3dClustSim are also shown in Fig. 1.

This mixed ACF model is now available in AFNI, but is not yet incorporated into the default processing stream (afni_proc.py).

For the per-voxel threshold of *p*=0.01, the impact of the bug fix is much smaller than that of the long tails in the mixed ACF model. For per-voxel threshold *p*=0.001, the impact of the bug fix is about the same as that of the long tails in the mixed ACF model (paired *t*-test), as these estimates were closer to the nominal 5% rate already. In every case, the bug fix reduced the FPR, as did the change to the mixed ACF model.

For the block design stimulus timings (“B1” and “B2”, 10 and 30 s blocks of “task”), the FPRs are still a little high even with the mixed ACF model. Preliminary results indicate this bias is partly due to the fact that the spatial smoothness of the FMRI noise is a function of temporal frequency-the FMRI noise at lower temporal frequencies is smoother than at higher frequencies. The two event-related stimulus timings (“E1” and “E2,” the regular and random 2 second events, respectively) are at higher temporal frequencies, so the noise that corrupt their results are somewhat less smooth than the noise corrupting the block design results. Since the smoothness parameters (FWHM or *a,b,c*) are estimated from *all* the residuals of the task-activation analysis, they tend to be biased towards the more numerous higher frequency bands.

### The Future-A Third Problem with Cluster-Threshold Detection Tools

A third standard assumption (present in AFNI, as well as FSL and SPM) also makes the idea of using a global cluster-size threshold somewhat questionable. In fact, the spatial smoothness of the FMRI noise is *not* spatially stationary-it is significantly smoother in some brain regions (eg, the precuneus and other large areas involved in the default mode network) than in others (this effect is also noted in [1,2]). Variable smoothness means that the density of false positives for a fixed cluster-size threshold will vary across the brain, especially since the FPR is strongly nonlinear in the cluster-size threshold. Using the same cluster-size threshold everywhere in such brain data can result in higher FPRs than expected in the smoother areas and lower FPRs than expected in the less-smooth areas.

One way to address this problem is to perform spatially variable smoothing, with the aim that each brain region reaches a target level of smoothness (as opposed to current approaches, which apply a single constant smoothing FWHM of a user-specified level). Since the ACF is not Gaussian-shaped, this approach requires making a local estimate of the (Eq 1) model parameters in order to guide the variable smoothing algorithm. A new AFNI program, 3dLocalACF, has been written to estimate the *a, b, c* parameters locally (in a ball, constrained within a brain mask) around each brain voxel. In this case, the non-Gaussian smoothness is characterized by a new parameter called the “Full-Width at Quarter-Maximum” (FWQM), which characterizes the scale of the model in (Eq 1) at a broader point than the FWHM used in the simple Gaussian case; in the limiting case that the ACF is Gaussian, then FWQM = 2^1/2^×FHWM.

An example of the FWHM and FWQM smoothness estimates for one subject are shown in Fig. 3. Work is underway to use this program’s results to control spatially variable smoothing to a target (via a finite difference approach to solving a parabolic PDE), and then test the FPR via simulation. Preliminary results are encouraging, but further simulations are needed before this method can be recommended for regular practice.

**Figure 3.**
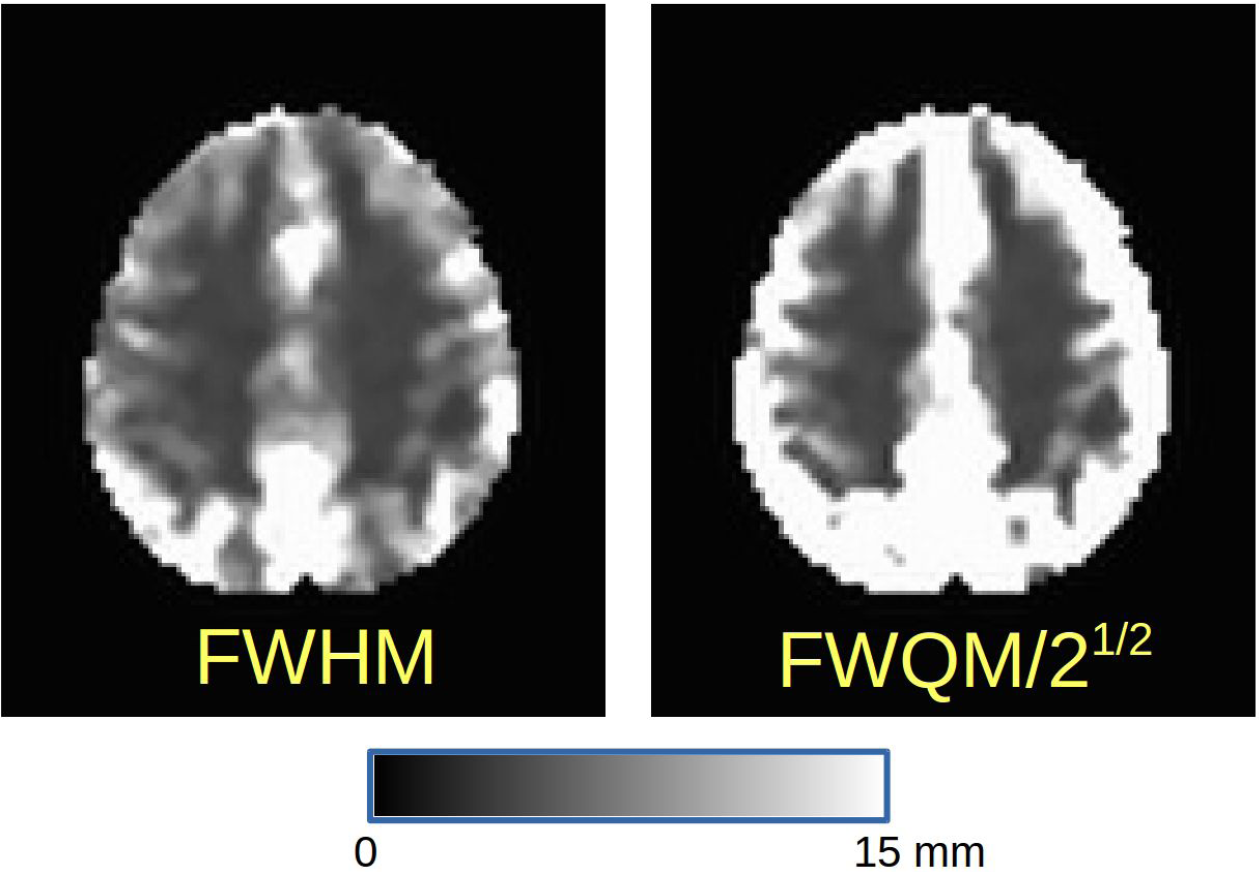
Images of the FMRI noise FWHM and the Full Width at Quarter Maximum (FWQM) from one subject (#11344) in the Beijing dataset (after nominal smoothing with a Gaussian kernel of 4 mm FWHM during preprocessing). The scale in both images is linear from black = 0 to white = 15 mm. If the ACF were Gaussian, FWQM = 2^1/2^×FHWM. The FWHM map shows that the noise smoothness is not uniform in space (even within gray matter), and the FWQM map shows that the non-Gaussianity of the noise smoothness is also non-uniform. The magnitude of this effect on the FPR and how to allow for it in thresholding are still under investigation.

### The Present-A Non-Parametric Approach to Cluster-Size Thresholding

A second approach to adjusting the FPR in cluster-size thresholding has been implemented in the AFNI program 3dttest++ (a program that is also capable of incorporating between-subjects factors and covariates, in addition to carrying out simple voxel-wise *t*-tests; this point is discussed more later). The procedure is straightforward:

- Compute the residuals of the model at each voxel at the group level;
- Generate a null distribution by randomizing among subjects the signs of the residuals in the test, repeat the *t*-tests (with covariates, if present), and iterate 10,000 times;
- Take the 10,000 3D *t*-statistic maps from the randomization and use those as input to 3dClustSim (with no additional smoothing): threshold the maps, cluster-ize them, and then count the false positives.

All these steps are carried out inside the 3dttest++ program, if the new command line option ‘-Clustsim’ is added. Each simulation run of 3dttest++ took 6-7 minutes of clock time to run on a 16 core node of the NIH Linux cluster (the randomized t-tests and 3dClustSim are multi-threaded).

The output is a table of cluster-size thresholds for a range of per-voxel *p*-value thresholds and a range of cluster-significance values. Such a table is produced for each of the clustering methods that AFNI supports: 1^st^, 2^nd^, and 3^rd^ nearest neighbors, and 1-sided or 2-sided voxel-wise thresholding. (In general, we prefer 2-sided *t*-statistic thresholding in AFNI as providing more transparency about and more rigorous FPR control of the results, but do allow for 1-sided thresholding.) These tables are saved in text format, and also stored in the header of the output statistics dataset for use in interactive thresholding in the AFNI GUI.

For comparison here, the 1000 2-sample *t*-tests described above were re-run for the 16 cases (4 blurring levels times 4 stimulus timings) with this new ‘-Clustsim’ option, and tested against each of the 6 combinations of thresholding-sidedness and clustering-neighborliness possible in AFNI, over a range of per-voxel p-value thresholds. The results were similar across all 96 cases. The results for the 1-sided 1^st^ nearest neighbor clustering approach are shown graphically in Fig. 4; all false positive rates are within the “nominal” 95% confidence interval for the FPR (3.65-6.35%) over the collection of per-voxel *p*-value thresholds tested. At this time, we recommend the use of this option for cases where the group analysis is simple enough to carry out via a GLM with 3dttest++ (such as a 1- or 2-sample *t*-test).

**Figure 4.**
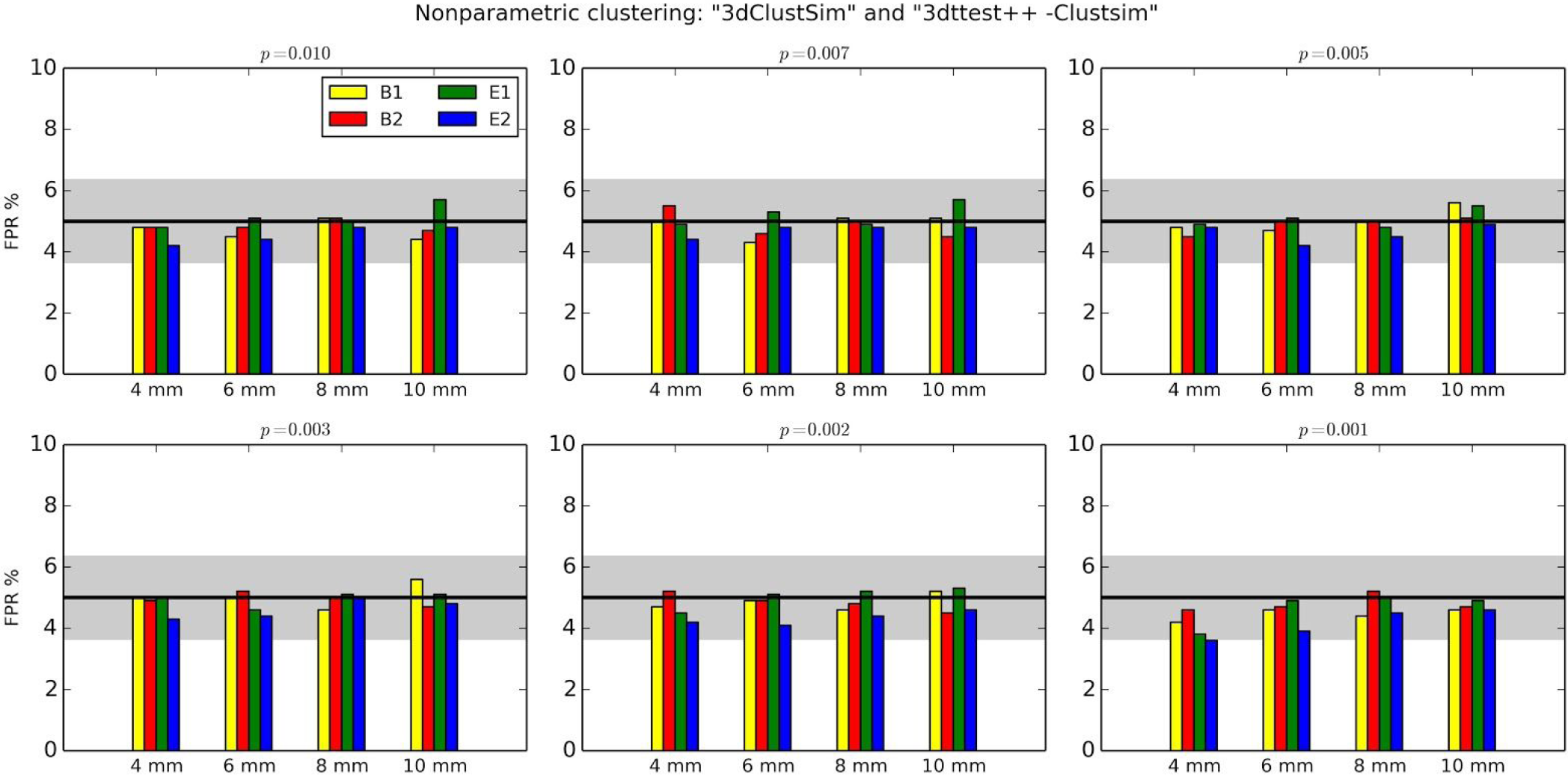
FPRs with cluster-size thresholds now determined from the ‘-Clustsim’ option of 3dttest++ (1-sided tests with 1^st^ nearest neighbor clustering). See Fig. 1 for description of labels, but note that the y-axis range has been changed here for visual clarity.

## Discussion and Conclusions

### A Note on “The Bug” and Bugs in General

Of the many features investigated here, we first comment on one of the most publicly highlighted ones-the bug in the older versions of 3dClustSim (and its precursor AlphaSim). As shown here and noted before, this is actually a minor feature-the effect of the bug on FPR is relatively small. Correcting the underlying feature did indeed reduce the false positive rates in these tests, but the change in results is not a major factor in the overall FPR inflation. Both before and after the bug fix, 3dClustSim performed in comparable manner to the other software tools being investigated [2]. This is not to say that the presence of the bug was not unfortunate, but it was not research-destroying. In order to have significant changes in the results of 3dClustSim, new methods were required, which were also presented and which are discussed further below.

“Reproducibility” has been a major topic in the field of FMRI, with several proposals of “best practices” put forth in various forms. It is obvious that the presence of bugs in software (as well as misuses of software settings inappropriate in the context) damages the validity and reproducibility of reported results, and there is no greater concern for those writing software-particularly when it is intended for public use-than preventing bugs. However, practical realities are that (a) not every user will program her/his own software tools, and (b) every large software distribution is essentially guaranteed to contain bugs (mistakes, invalid assumptions, etc) at some level. Much of the discussion surrounding [2], particularly in comments to and take-aways chosen by the popular press, focused on the bug that was present in 3dClustSim. The discovery of this bug was highlighted and hyped as a major component for rejecting 15 years of brain studies and (up to) 40,000 peer-reviewed publications on the brain^1^, under the tacit or explicit assumption that the reported results would be unreproducible.

However, rather than being evidence for “a crisis of reproducibility” within the field of FMRI, the advertisement of the bug is itself an important verification of the reproducibility of FMRI analysis. In this imperfect-world context, the philosophy for maintaining AFNI has always been to correct any bugs and to update the publicly available software as soon as practicable, often posting on the public Message Board for significant changes, and maintaining a permanent, public list of updates/changes/bugs online. This specific bug, which had been tested and found to be small, was able to be advertised so publicly in part because the software maintainers actively advertised it (see the AFNI change log note in the Introduction).

While certainly an annoying moment for both the researchers who used 3dClustSim and those who maintain the software, the knowledge and dissemination of this bug is part of the reproducibility process. The existence of software bugs is unfortunate but likely inevitable-even huge distributions such as Python, Windows, Mac, and Linux release bug fixes regularly. Clarity of description and speed of repair are the best tools for combatting their effects once discovered.

### The state of clustering

There were many valuable points raised in the work of [2]. Several of these were important for general consideration within the FMRI field, such as the assumption of most clustering approaches that spatial smoothness was well-enough approximated by a Gaussian shape. To address this point, we have shown how an updated approach within AFNI using an estimated non-Gaussian ACF greatly improves the FPR controllability within the test datasets. Additionally, there is also a new nonparametric method for clustering within AFNI that shows promise; however, this type of approach currently appears to be limited by practical considerations (that hold across software implementations) to relatively basic group analyses that can be performed through univariate GLM.

Further work will be required to more fully develop satisfactory cluster detection levels within FMRI, in particular being able to address the inhomogeneity of smoothness and structure in the brain while trying to detect ‘true positives’ that are either in very small regions (eg, the hippocampus and amygdala) or very large volumes. In the absence of a “gold standard” correct answer, further methods development will continue, likely without producing a single “best” answer. For example, the authors of [2] investigated 4 different software packages, and there were 6 different “standard” methods among them! While computer power is impressive these days, very few labs perform daily analyses on GPUs, and even so, permutation tests of complex models (e.g., complex AN(C)OVA or LME) become extremely computationally expensive. Furthermore, issues of smoothness and the inhomogeneity of both underlying and noise structure in brain images present challenges for any method.

An interesting feature of the large volume of work performed by the authors of [2] is that there were consistent differences in method performance based on simulation paradigm and parameters themselves. This suggests that a number of issues with FPRs can also be addressed significantly with careful experimental design-for example, the benefits of event-related designs and, in particular, random event-related tasks were apparent in all cases. The choice of voxelwise *p*-value is also paramount-*all* software packages produced fairly reasonable results for *p*~0.001 or below; in fact, this single user-defined choice could solve many of the problems presented by [2] to a very large degree.

### Is a Parametric Approach to Cluster-Size Thresholding Desirable?

A permutation or randomization approach seems able to provide proper FPR control with few apparent assumptions; so why not use this approach for everything in FMRI group analysis?

The primary answer is practical: not all statistics are easily re-computable thousands of times. For example, linear mixed effects (LME) analysis requires nonlinear regression at each voxel, and is computationally intensive itself [10]. Repeating it massively with permuted or randomized data is not feasible at the present juncture.

A secondary answer is that it is not easy nor practicable to permute or randomize in complicated group analysis setups, with a mixture of between- and within-subject factors and covariates, multi-way AN(C)OVA-like analyses outputting various statistics to test for interactions, conjunctions, etc. Additionally, a permutation test is not always required, and it may sacrifice power in some cases (e.g., a fixed number of permutations would set a lower bound for the *p*-value that could be achieved, leading to failure to detect a small cluster with a potentially very high significance level that would survive through a parametric approach). While permutation testing may be useful and even necessary in some situations, a general rule for determining those cases is not clear, and it may be computationally or methodologically prohibitive to use (e.g., in the common case of including covariates, mixed effects, etc.). Further work is required on this important issue.

### Final (for now) Thoughts on Statistics in FMRI

While false positive rates are important, we cannot also forget about not wanting to use methods that give overly large false *negative* rates. The presence of persistent over-stringency in clustering methods (eg, failing to survive statistical significance thresholding when real effects actually exist) would be as much a problem for interpreting findings in the brain as highlighting weak differences. Certainly, when using a cluster-wise *p*-value, one would hope that a method would reliably reflect the nominal rates. But, in conjunction with other trending discussions in the statistics literature, p-value thresholds are not sacred boundaries so that results around them live or die by tiny fractions above or below them. Thresholding is a convenience for focusing reporting, but they are only part of the story. Our point here ties into discussions of reducing ‘p-hacking’ and emphasizing effect sizes in results reporting for FMRI [11].

Statistical testing and reporting is far from the end of a neuroscientific FMRI paper; in fact, it is just the technical prelude to the neuroscientific interpretation-which is not (often) statistical at the present time, but rather is qualitative, built on previous work and knowledge, adding in new information from the present experiment. It is *very hard* to decide without close examination if a weak positive cluster in a brain map is actually critical to forming the paper’s conclusions.

In other words, don’t throw the baby out with the bathwater-just state clearly if “the baby” is borderline statistically significant.

## Acknowledgements

The research and writing of the paper were supported by the NIMH and NINDS Intramural Research Programs (ZICMH002888) of the NIH/DHHS, USA. This work utilized the computational resources of the NIH HPC Biowulf cluster (http://hpc.nih.gov).

## Supplementary Tables: Data used in Figures 1 and 4

**Supplementary Table 1:** FPRs for various software scenarios, with 1000 2-sample *t*-tests (as in [1,2]) using 20 subjects’ data in each sample. “buggy” and “fixed” means the cluster-size thresholds selected using the Gaussian shape model with the FWHM being the median of the 40 individual subject’s values; “buggy” is using 3dClustSim before the bug fix, “fixed” is using 3dClustSim after the bug fix, “mixed ACF” means the cluster-size threshold selected using the (Eq 1) model for spatial correlation of the noise, with the *a,b,c* parameters being the median of the 40 individual subject’s values (estimated via program 3dFWHMx). Two different per-voxel *p*-value thresholds (1-sided tests, as used in [2]) are shown. The 95% confidence interval for the expected 5% false positives out of 1000 trials is 0.036-0.064.

|  |  | buggy | fixed | mixed ACF | buggy | fixed | mixed ACF |
| --- | --- | --- | --- | --- | --- | --- | --- |
| blur | stim | $p=0.01$ | $p=0.01$ | $p=0.01$ | $p=0.001$ | $p=0.001$ | $p=0.001$ |
| 4 mm | blk10 (B1) | 0.384 | 0.355 | 0.274 | 0.117 | 0.111 | 0.109 |
| 6 mm | blk10 | 0.406 | 0.358 | 0.203 | 0.149 | 0.123 | 0.097 |
| 8 mm | blk10 | 0.367 | 0.346 | 0.276 | 0.125 | 0.114 | 0.113 |
| 10 mm | blk10 | 0.321 | 0.272 | 0.137 | 0.125 | 0.108 | 0.087 |
| 4 mm | blk30 (B2) | 0.367 | 0.346 | 0.276 | 0.125 | 0.114 | 0.113 |
| 6 mm | blk30 | 0.352 | 0.322 | 0.192 | 0.124 | 0.109 | 0.098 |
| 8 mm | blk30 | 0.317 | 0.260 | 0.162 | 0.112 | 0.096 | 0.081 |
| 10 mm | blk30 | 0.250 | 0.222 | 0.136 | 0.097 | 0.083 | 0.064 |
| 4 mm | ereg (E1) | 0.150 | 0.136 | 0.071 | 0.033 | 0.030 | 0.028 |
| 6 mm | ereg | 0.217 | 0.173 | 0.071 | 0.079 | 0.064 | 0.042 |
| 8 mm | ereg | 0.242 | 0.178 | 0.078 | 0.100 | 0.071 | 0.048 |
| 10 mm | ereg | 0.231 | 0.181 | 0.078 | 0.106 | 0.075 | 0.049 |
| 4 mm | eran (E2) | 0.238 | 0.212 | 0.125 | 0.069 | 0.062 | 0.057 |
| 6 mm | eran | 0.257 | 0.240 | 0.101 | 0.099 | 0.074 | 0.059 |
| 8 mm | eran | 0.288 | 0.232 | 0.097 | 0.103 | 0.075 | 0.059 |
| 10 mm | eran | 0.259 | 0.215 | 0.092 | 0.100 | 0.078 | 0.059 |

**Supplementary Table 2:**
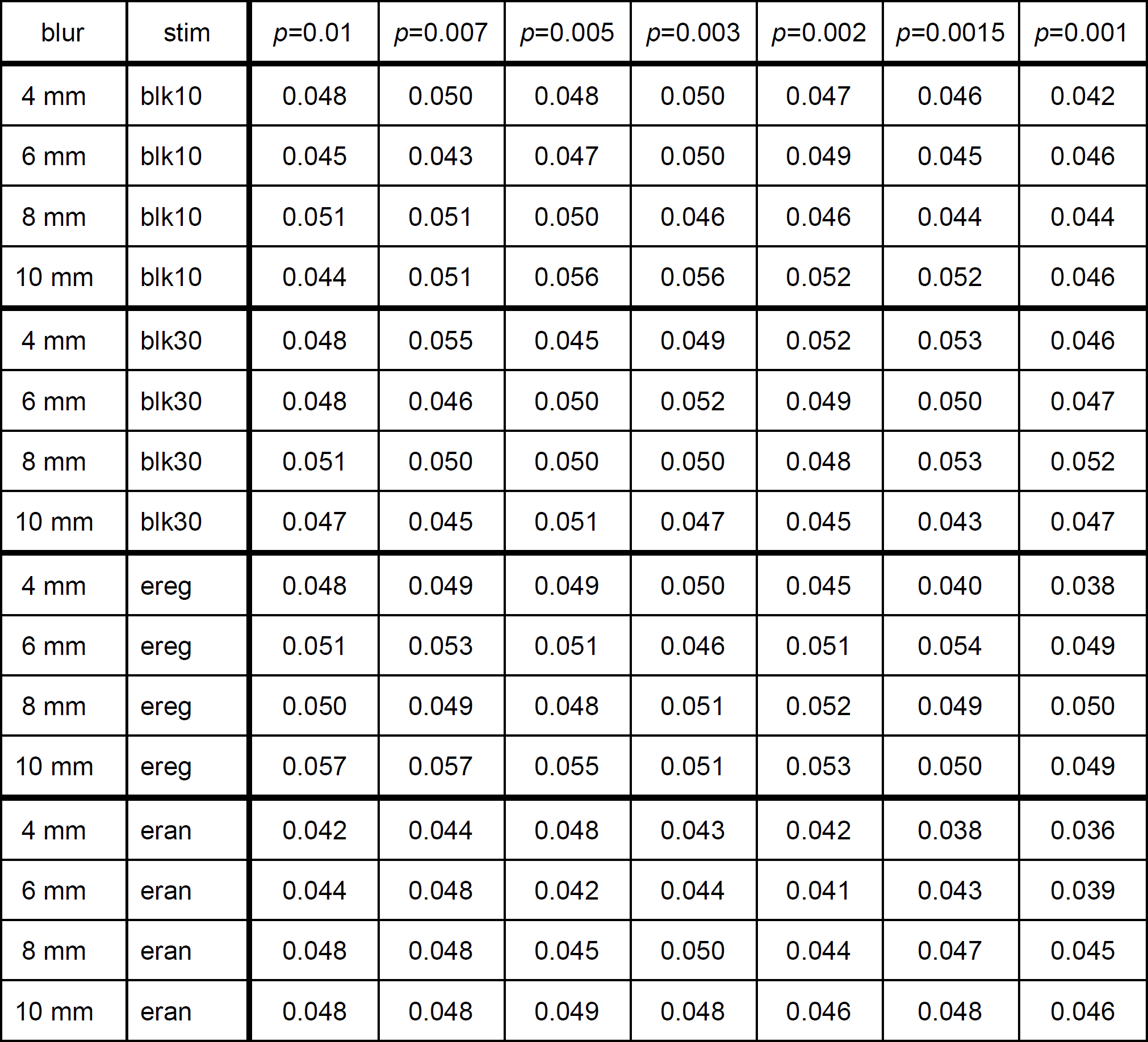
Analogous to Table 1, but with cluster-size thresholds now determined from the ‘-Clustsim’ option of 3dttest++ (1-sided tests with 1^st^ nearest neighbor clustering). Results from the 5 other possible combination of sidedness and neighborliness are very similar.

| blur | stim | $p=0.01$ | $p=0.007$ | $p=0.005$ | $p=0.003$ | $p=0.002$ | $p=0.0015$ | $p=0.001$ |
| --- | --- | --- | --- | --- | --- | --- | --- | --- |
| 4 mm | blk10 | 0.048 | 0.050 | 0.048 | 0.050 | 0.047 | 0.046 | 0.042 |
| 6 mm | blk10 | 0.045 | 0.043 | 0.047 | 0.050 | 0.049 | 0.045 | 0.046 |
| 8 mm | blk10 | 0.051 | 0.051 | 0.050 | 0.046 | 0.046 | 0.044 | 0.044 |
| 10 mm | blk10 | 0.044 | 0.051 | 0.056 | 0.056 | 0.052 | 0.052 | 0.046 |
| 4 mm | blk30 | 0.048 | 0.055 | 0.045 | 0.049 | 0.052 | 0.053 | 0.046 |
| 6 mm | blk30 | 0.048 | 0.046 | 0.050 | 0.052 | 0.049 | 0.050 | 0.047 |
| 8 mm | blk30 | 0.051 | 0.050 | 0.050 | 0.050 | 0.048 | 0.053 | 0.052 |
| 10 mm | blk30 | 0.047 | 0.045 | 0.051 | 0.047 | 0.045 | 0.043 | 0.047 |
| 4 mm | ereg | 0.048 | 0.049 | 0.049 | 0.050 | 0.045 | 0.040 | 0.038 |
| 6 mm | ereg | 0.051 | 0.053 | 0.051 | 0.046 | 0.051 | 0.054 | 0.049 |
| 8 mm | ereg | 0.050 | 0.049 | 0.048 | 0.051 | 0.052 | 0.049 | 0.050 |
| 10 mm | ereg | 0.057 | 0.057 | 0.055 | 0.051 | 0.053 | 0.050 | 0.049 |
| 4 mm | eran | 0.042 | 0.044 | 0.048 | 0.043 | 0.042 | 0.038 | 0.036 |
| 6 mm | eran | 0.044 | 0.048 | 0.042 | 0.044 | 0.041 | 0.043 | 0.039 |
| 8 mm | eran | 0.048 | 0.048 | 0.045 | 0.050 | 0.044 | 0.047 | 0.045 |
| 10 mm | eran | 0.048 | 0.048 | 0.049 | 0.048 | 0.046 | 0.048 | 0.046 |

## Footnotes

1 These estimates have since been greatly reduced and de-emphasized by the authors. In a paper focused on reducing the number of false positives in FMRI studies, it is slightly ironic that there were such large “false positive” conclusions reported.

